# Clear optically matched panoramic access channel technique (COMPACT) for large-volume deep brain imaging

**DOI:** 10.1101/2020.09.16.300103

**Authors:** Bowen Wei, Chenmao Wang, Zongyue Cheng, Wenbiao Gan, Meng Cui

## Abstract

To understand neural circuit mechanisms underlying behavior, it is crucial to observe the dynamics of neuronal structure and function in different parts of the brain. Current imaging technologies allow cellular resolution imaging of neurons within ~1 millimeter below the cortical surface. Even for mice, the majority of brain tissue remains inaccessible. Although miniature optical imaging probes allow the access of deep brain regions, the cellular-resolution imaging is restricted to a small tissue volume. To drastically increase the tissue access volume and enable a high-throughput neurophotonic interface, we developed a clear optically matched panoramic access channel technique (COMPACT). With comparable probe dimension, COMPACT enables a two to three orders of magnitude greater tissue access volume for structural and function imaging. Leveraging the large-volume imaging capability of COMPACT, we demonstrated multiregional calcium imaging of deep brain functions associated with sleep for the first time. The compatibility of COMPACT for longitudinal large-volume *in vivo* imaging will be highly valuable to a variety of deep tissue imaging applications.

---

High resolution optical imaging in the living brain has become a powerful tool for investigating the plasticity and function of neural circuits underlying animals’ behaviors.^1–8^ The latest advances in genetically encoded fluorescent indicators and optical imaging have enabled selective labeling and observation of neuronal structure and function in live animals, which have profoundly transformed the study and understanding of neural circuits. Such powerful technologies rely on our capability of delivering focused light inside brain tissue. Due to the refractive index inhomogeneity induced random light scattering, single cell resolution function imaging is limited to ~1 millimeter (mm).^9–11^ Even for the centimeter scale mouse brain, such penetration depth can only access superficial brain regions. The majority of the brain volume remains inaccessible to high-resolution optical imaging. Although macroscale and mesoscope imaging modalities such as fMRI and ultrasound based methods allow the imaging of deep brain structures, they lack the single cell resolution and sensitivity which is crucial to the understanding of neural circuitry.^12^ The current method of choice for cellular resolution deep brain imaging requires inserting miniature optical probes.^13–21^ However, a major limitation of such practice is the very limited tissue access volume which severely limits both the throughput and the success rate of practical implementation. If the neurons of interest were not located within the probe’s small imaging volume, new animals and surgeries would be needed. New techniques that can provide orders of magnitude more tissue access volume for longitudinal high-resolution imaging and thus greatly improve the throughput, flexibility and success rate are highly desirable.

Several invasive approaches have been developed for optical recording of deep brain structures, such as the excision of overlying brain tissue, miniature prism implantation, miniature gradient-index (GRIN) lens probes and their combinations.^13, 15–20^ For viewing very deep brain regions, miniature lens probes are typically employed due to the minimized tissue damage. Such miniature lens probes typically utilize GRIN lens for its slim cylindrical body to form focus deep inside brain tissue and relay the optical signal to the outside for detection. The field of view (FOV) of GRIN lens utilized for deep brain two-photon imaging is typically ~1/5 of the lens diameter. Greater lens diameter can provide greater FOV at the cost of more tissue damage. However, the ratio of imaging volume to insertion volume barely changes. New innovations are needed to overcome the access volume and scale limitation.

Here we report the development of a clear optically matched panoramic access channel technique (COMPACT), which drastically overcomes the tissue access volume constraint. With the same insertion device volume, COMPACT can provide two to three orders of magnitude increase in tissue access volume. Specifically, COMPACT enables a 360-degree panoramic view around the inserted probe throughout the probe length. As such, the entire probe surface is accessible for high-resolution imaging. The key idea of COMPACT is to abandon the convention of directly inserting imaging lens inside brain tissue. Instead, we insert a clear optically matched channel inside the tissue. To form images, we then insert an imaging probe with side viewing capability inside the channel, similar to the side-view endoscope used in medical applications.^22, 23^ With the channel isolating the imaging probe from the tissue and providing refractive index matching with the tissue, we could freely spin and translate the imaging probe inside the channel to image different brain regions. The spinning of the imaging probe enables a 360 degree panoramic view around the channel. The translation of the probe along the channel allows us to image through the entire insertion length. Compared to the tiny access volume restricted to the tip of conventional imaging probes, COMPACT allows large volume imaging all around the inserted device.

The design of the COMPACT system for *in vivo* two-photon imaging of head-fixed mice (**Fig. 1a**) is similar to that of conventional two-photon microscopes equipped with a spatial light modulator (SLM) (**Supplementary Video 1**). We imaged the two-axis galvo scanner onto the SLM which provided aberration correction and focal depth adjustment. The SLM was subsequently imaged onto the back focal plane of an objective lens whose focus was relayed to the brain tissue by a GRIN lens mounted on a precision rotary stage. To provide the side-viewing capability, we attached a high refractive index right-angle micro prism onto the facet of the GRIN lens.

**Figure 1.**
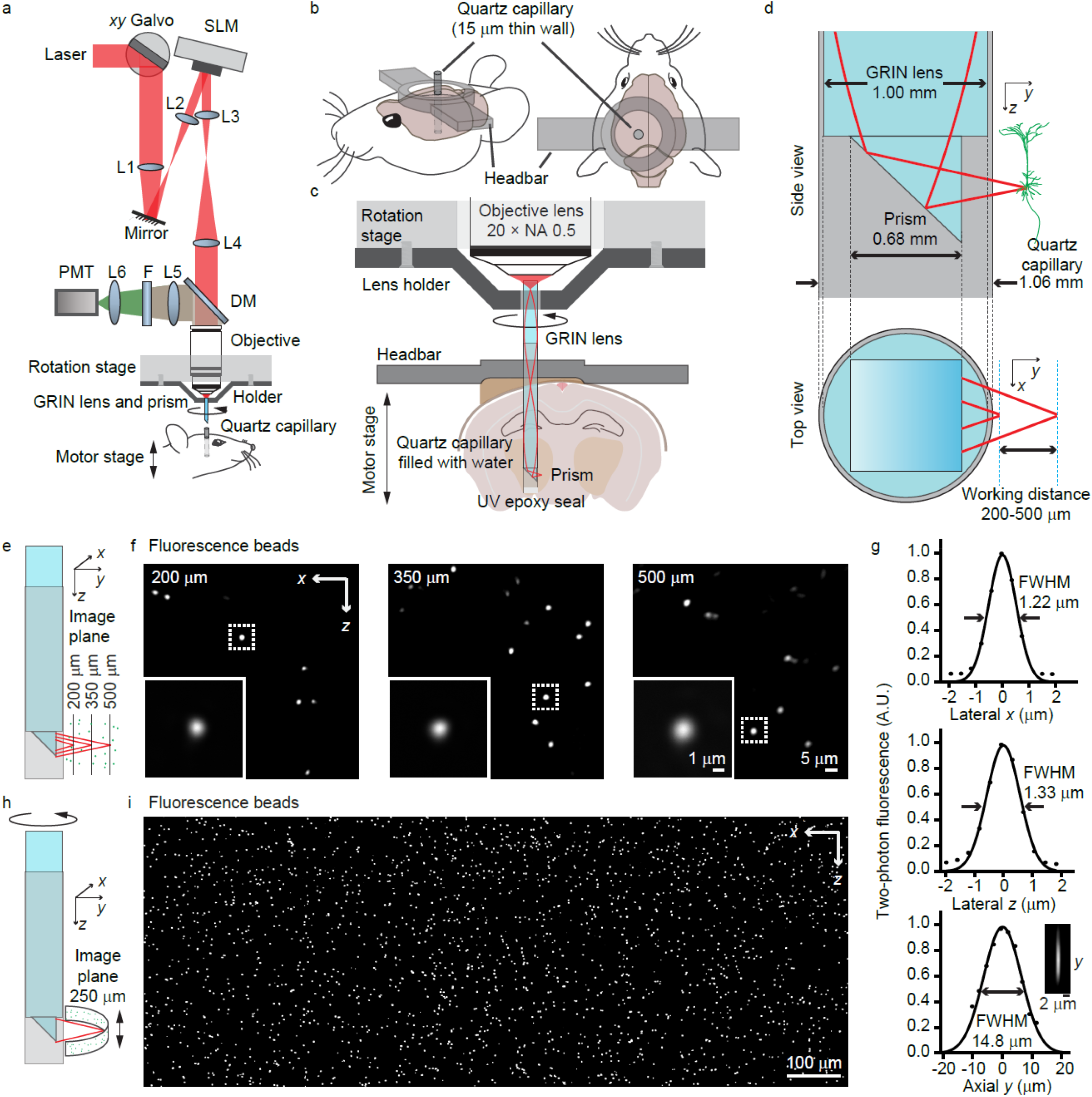
System design of COMPACT. (a) The optical design for COMPACT based two-photon imaging system. L1-4, telecentric relay lenses; L5-6, high etendue fluorescence collection lenses; F, bandpass filter; DM, dichroic mirror. (b) The configuration of head-bar and the surgically implanted quartz capillary. (c) The rotation and translation control for whole-depth panoramic imaging. The rotary stage spins the imaging probe for panoramic view and the motorized stage moves the capillary along the rotation axis for viewing different depths in the brain. (d) Dimension of the capillary, imaging probe and the working distance. (e) For PSF quantification, we parked the imaging probe and used SLM to move the laser focus to image beads at different distances from the capillary. (f) Images of 0.5 μm fluorescence beads. (g) The cross-section plot of the PSF at 200 μm outside the prism surface. (h) For panoramic large FOV imaging, we rotated and translated the imaging probe with respect to the sample. (i) Example of large FOV panoramic imaging of 0.5 μm fluorescence beads in agar. The range of angle rotation is 120 degree and the range of axial translation is 510 μm.

To form the clear optically matched channel, we surgically inserted an ultrathin quartz capillary with a wall thickness of 15 μm (**Fig. 1b**). We filled the capillary with water to match the refractive index of brain tissue and then inserted the imaging probe inside the capillary for imaging (**Fig. 1c**). It is worth noting that the 15 μm thin quartz induced aberration for two-photon excitation with NA ≤ 0.5 is insignificant. Moreover, the SLM was employed to compensate the overall system aberration in the imaging path, which was dominated by the GRIN lens induced spherical aberration.

For panoramic imaging around the capillary, we need to precisely spin the imaging probe around the rotation axis of the rotary stage. Thus, both the probe and the capillary need to be well aligned with respect to the rotation axis. Utilizing three machine vision cameras, we implemented an automatic alignment procedure (**Supplementary Fig.1**). These cameras also guided the process of inserting the imaging probe inside the channel. After the insertion, we could freely spin the imaging probe for panoramic imaging by controlling the rotary stage. Moving the motorized stage that held the mouse headbar along the rotation axis allowed us to image through the entire length of the capillary.

With the SLM controlled defocusing, we could typically image ~500 μm outside the prism facet, *i.e*. ~ 300 μm into the brain tissue (**Fig. 1d**). To calibrate the imaging performance, we embedded 0.5 μm beads in agar as the sample, inserted the quartz capillary inside the agar and then performed volumetric imaging (**Fig. 1e-1i**). With the simple probe design that only utilized a GRIN lens and a high index right-angle prism, the achieved spatial resolution (**Fig. 1g**) was sufficient for resolving the dendrites and somata of neurons. The excitation numerical aperture (NA) for imaging 200-500 μm outside the prism facet was 0.38-0.32 while the signal collection NA was ~0.5, limited by the objective lens. It is interesting to note that just slight rotation (120 degree) can yield very wide FOV (**Fig. 1h** and **1i**). For example, spinning the 1 mm diameter probe over 360 degrees while imaging ~300 μm outside the channel can yield a ~5 mm wide panoramic view. Such a FOV is impossible to obtain with conventional probes.

To evaluate COMPACT’s performance for *in vivo* structural imaging, we imaged the brain of eGFP mice at various depths and regions (**Fig. 2, Supplementary Fig. 2** and **Video 2**). With the flexibility of COMPACT, we have the freedom to visit a large volume of brain tissue. Experimentally, we defined the imaging probe’s insertion range and spinning range of interest in the control program, which automatically positioned the imaging probe to visit the defined locations in sequence. At each location, the SLM driven defocusing synchronously coordinated with the galvo scanning to record 3D image stacks. As an example, we show the millimeter scale images acquired from the neocortex and the hippocampus (**Fig. 2b-2d**, **Supplementary Fig 2**). Even with the moderate NA employed in this study, we could observe fine neuronal structures.

**Figure 2.**
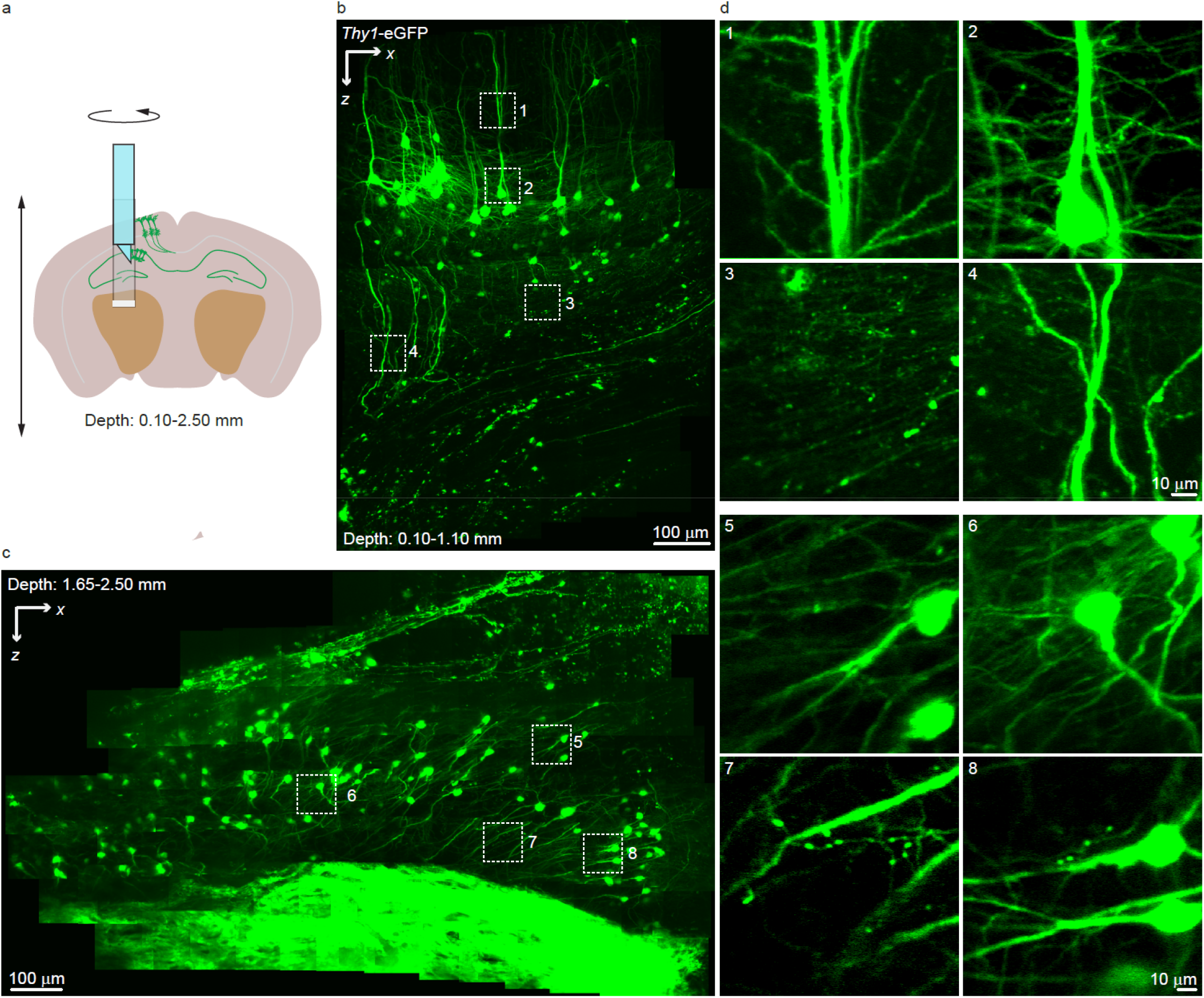
*In vivo* imaging of neuronal structure at different depths in eGFP mouse brain. (a) The spatial configuration for the panoramic imaging from the mouse cortex to hippocampus. (b) *In vivo* imaging of neurons from 0.10 to 1.10 mm in the mouse brain. (c) *In vivo* imaging of neurons from 1.65 to 2.50 mm in the mouse brain. (d) Zoomed-in view shows the fine features of neurons. The data were recorded at 1.7 Hz frame rate with ~20 mW laser power. The images shown are the average of five frames.

To evaluate COMPACT’s performance for *in vivo* calcium imaging, we imaged the GCaMP6s expressing neurons in the brain of awake mice (**Fig. 3, Supplementary Video 3**). Following the virus injection, we surgically inserted the quartz capillary (**Supplementary Fig. 3**). Four weeks after the surgery, we carried out *in vivo* imaging in head-fixed awake mice. As an example (**Fig. 3b–3f**), we show the recorded spontaneous calcium transients deep in hippocampus. With the two-photon based high-contrast imaging, we captured the calcium signal with high signal to noise ratio (SNR) from not only the somata (noise ~6% Δ*F*/*F*_0_) but also the dendrites (noise ~10% Δ*F*/*F*_0_) with large FOV in deep brain. It is worth noting that the COMPACT method can handle substantial tissue motion during awake animal calcium recording. Utilizing a simple imaging probe docking method (**Supplementary Fig. 4**), we allowed the imaging probe to flexibly move with the quartz capillary and therefore were able to tolerate significant tissue movement (**Supplementary Video 4**). These imaging results from awake mice indicate that the COMPACT system is well compatible with *in vivo* calcium signal recording.

**Figure 3.**
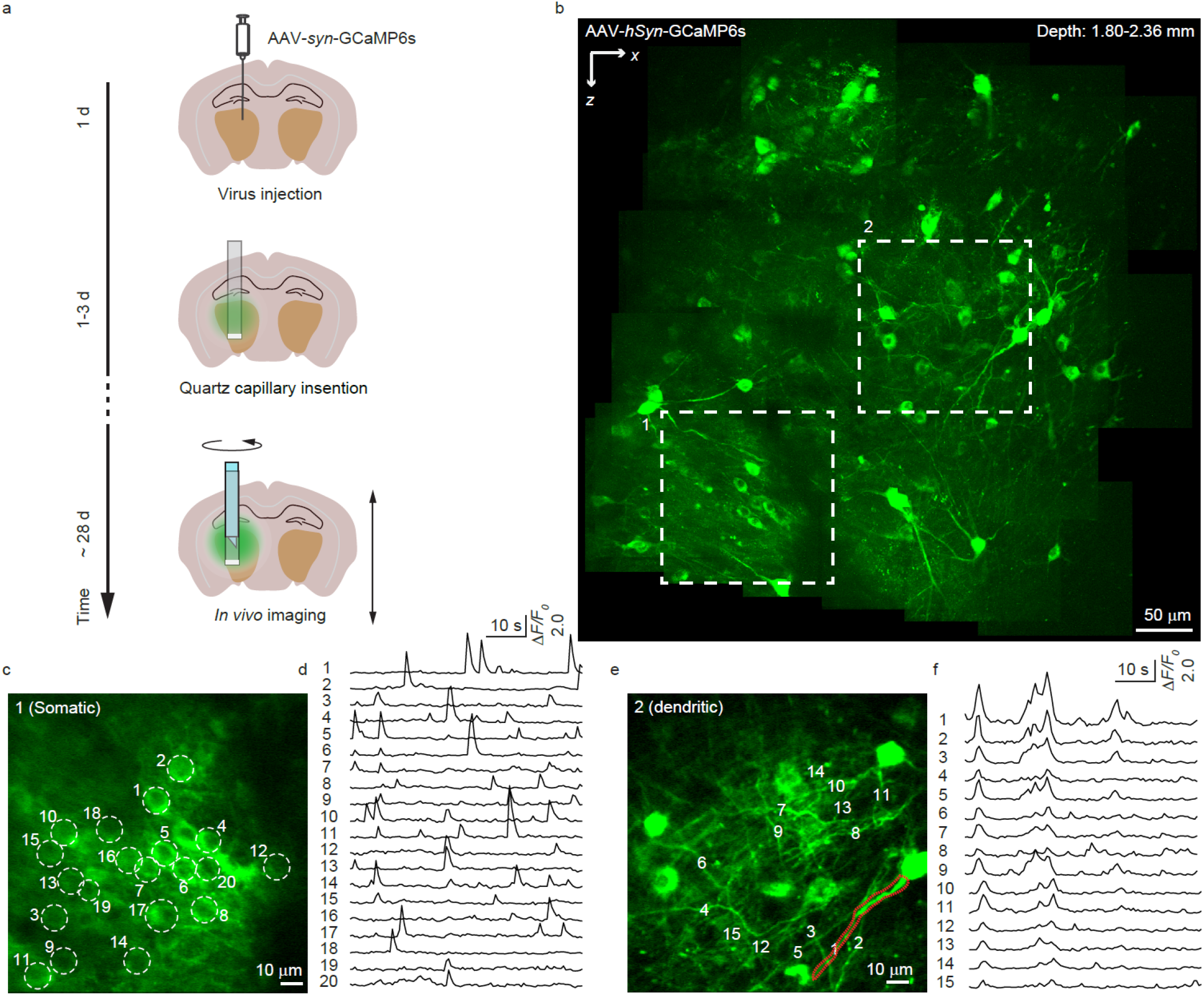
*In vivo* calcium imaging in the brain of a head-fixed awake mouse. (a) The timeline for virus injection, capillary implantation and *in vivo* imaging. (b) Structural imaging of GCaMP6s expressing neurons from 1.80 to 2.36 mm in the mouse brain. (c, d) Representative calcium transients from somas. (e, f) Representative calcium transients from dendrites. The images were recorded at 1.7 Hz frame rate with ~28 mW laser power.

As mammalian behaviors often involve neuronal population from multiple function regions, it is important to survey the neuronal activity over large brain volume. Leveraging the large volume accessibility of COMPACT, we performed calcium imaging across multiple regions associated with sleep (**Fig. 4**, **Supplementary Video 5**). Experimentally, we injected GCaMP6s virus (2 mm posterior and 2 mm lateral from Bregma) and implanted quartz capillary. After 4 weeks of recovery, we used 5-7 days to habituate the mice to sleep under head restraint condition. We then employed the two-photon COMPACT system to record the neuronal calcium activity while monitoring the pupil size to assess the stage of sleep^24^. Each recording session lasted for 20 minutes. From the correlation between pupil area and calcium transient (**Fig. 4f** and **4h**), we found that the neurons in the posterior thalamic nuclear (ROI 1 in **Fig. 4**) were highly active during sleep (pupil smaller than 0.15 mm^2^) and were quiet while awake (pupil greater than 0.4 mm^2^). Statistics showed that the neuronal activity and the pupil area had a strong negative correlation (**Fig. 4j** and **4k**). In comparison, we translated the imaging probe upward by ~ 1 mm and recorded the calcium transients from neurons in dentate gyrus (ROI 2 in **Fig. 4**). There were slight calcium activities during both asleep and awake states. However, during the transition from sleep to awakening, the neuronal populations showed highly synchronized calcium activity (**Fig. 4g** and **4i**). The statistics (**Fig. 4l** and **4m**) showed no direction correlation as that of the neurons in ROI1. As the COMPACT system offers high repeatability (**Supplementary Fig. 5**), we observed the calcium activity of ROI 1 and 2 neurons for two consecutive days with four imaging sessions per day. Similar thalamic region imaging experiments were repeated on six mice. Overall, the repeated longitudinal recording over different function regions demonstrated in this study highlights the multiregional imaging capability of COMPACT.

**Figure 4.**
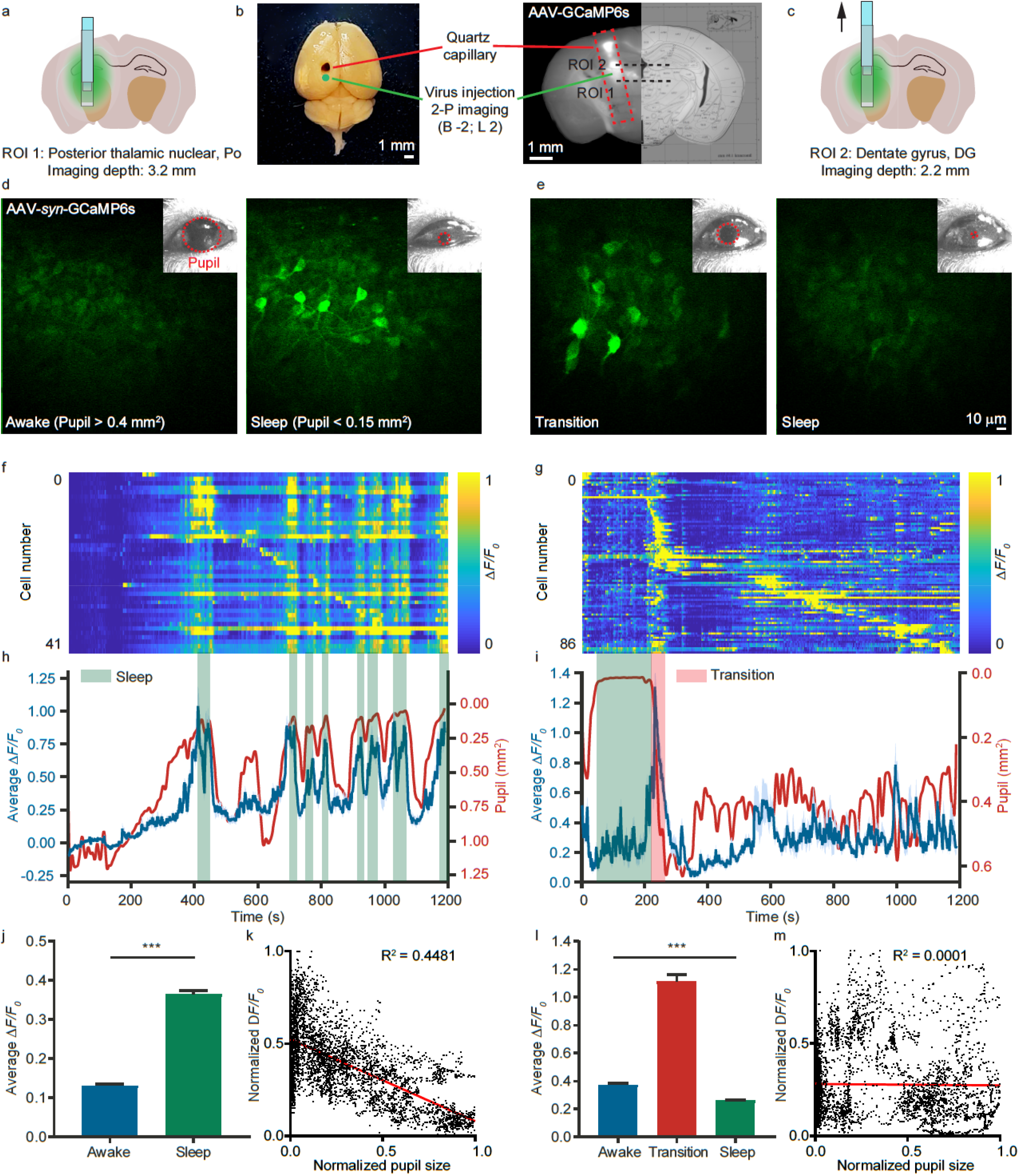
Neurons in different function regions showed distinct activity associated with sleep. (a) Position of ROI 1 (posterior thalamic nuclear). The depth is 3.2 mm. (b) Images of the dissection. Scale bar: 1 mm. (c) Position of ROI 2 (dentate gyrus). The depth is 2.2 mm. (d, e) Representative images showing the calcium activity of neurons during awake and asleep states in ROI 1 and 2, respectively. The images were recorded at 0.84 Hz frame rate with ~28 mW laser power. The insets show the images of the pupil recorded simultaneously with calcium imaging. Scale bar: 10 μm. (f, g) The calcium transients of individual neurons over 20 minutes sorted by firing time in ROI 1 and 2, respectively. (h, i) The average *ΔF/F_0_* (blue) and the pupil area (red) in ROI 1 and 2, respectively. The green boxes indicate the time windows when the pupil area was smaller than 0.15 mm^2^, and the red box indicates the time window of transitioning from sleep to awakening. (j) The average calcium transients of neurons in ROI 1 during awake and asleep states (n = 159, 138). (k) Correlation analysis between calcium transient in ROI 1 and pupil area (n = 4000; total recording time, 80 mins). (l) The average calcium transients of neurons in ROI 2 during the awake, transition, and asleep states (n = 191, 155, 256). (m) Correlation analysis between calcium transients in ROI 2 and pupil area (n = 4000; total recording time, 80 mins).

Compared to common miniature imaging probe based calcium recording, COMPACT has its key advantages in its massive tissue access volume and adaptability. Take the 1 mm diameter GRIN lens probe as an example, the conventional method can typically deliver a ~0.2 mm diameter two-photon imaging FOV and image ~0.3 mm outside the device, yielding a tissue access volume of 0.009 mm^3^. In comparison, COMPACT of the same insertion diameter can allow a circular imaging FOV of ~5 mm. Assuming that the translation along the capillary is 3 mm, COMPACT offers a tissue access volume of ~4 mm^3^, which is near three orders of magnitude greater than that of conventional methods. Such a huge increase in access volume provides tremendous flexibility to explore the neurons of interest and therefore greatly increases the success rate of finding neurons relevant for the studies and the overall experiment throughput. In addition, we also have the freedom to switch the imaging probe to visit the same population of neurons. For example, we could switch between probes of different NA (**Supplementary Fig. 6**) for the tradeoff between imaging volume and spatial resolution, similar to switching objective lenses under a conventional microscope, or switch between one photon wide field recording and two-photon imaging for the tradeoff between imaging throughput and imaging depth. One additional benefit of COMPACT is that we will no longer need many imaging probes, one for each animal. Instead, a high-quality and high-performance imaging probe can be utilized to imaging many animals as only quartz capillaries are inserted inside the brain tissue.

In summary, we developed COMPACT for large-volume neurophotonic interface. Instead of implanting imaging probes directly inside the brain tissue, we insert a clear and optically matched channel inside brain tissue, which allows us to freely translate and spin the imaging probe inside the brain to obtain a 360-degree panoramic view through the entire capillary insertion length. In fact, with the device’s entire surface assessable for high-resolution imaging, the design of COMPACT has maximized the ratio of imaging volume to device volume. Experimentally, we employed this system for two-photon imaging of neuronal structures and calcium activity in the brain of head-fixed awake mice. The high-SNR and high-contrast images indicate that COMPACT is compatible with routine calcium recording. The longitudinal multiregional imaging of neuronal activity associated with sleep further highlights the flexibility and volume advantages of COMPACT. Further engineering in miniaturization and imaging speed is expected to enable the COMPACT based whole mammalian brain access neurophotonic interface for many animal models.

## Methods

### Motorized capillary insertion

One key step to a successful implementation of miniature probe based deep brain imaging is the surgical insertion procedure. To make COMPACT based imaging robust and reproducible, we developed a motorized capillary insertion procedure (**Supplementary Fig. 3**). With the goal of imaging head-fixed animals, we first attached the head-bar to the skull. Using a dental drill, we carefully removed the skull near the middle of the head-bar above the insertion spot. Next, we used a computer controlled motorized actuator (Z825, Thorlabs) to gradually insert an open-end 15 μm thin quartz capillary into the brain tissue at 10 μm/sec insertion speed. It is worth noting that the 15 μm thin quartz wall is actually thinner than the common razor edge, which allows it to easily glide inside the tissue. During this process, we employed a 3-axis precision linear stage (461-XYZ-M, Newport) to position a thin needle (21G, BD) connected to a vacuum pump to remove the tissue inside the capillary. After the capillary reached the desired depth and the removal of the tissue inside the capillary, we reversed the motion of the motorized actuator that held the capillary and moved it out of the brain at 10 μm/sec speed. Next, we loaded a new 15 μm thin quartz capillary whose bottom was sealed with a tiny amount of epoxy inside the injection guiding chamber. The chamber was formed by the glass capillary and piston rod utilized inside transfer pipettes. Again, we utilized a motorized actuator to push the piston down at 10 μm/sec speed. During the whole insertion process, we constantly flushed the area with artificial cerebrospinal fluid (ACSF) to avoid bleeding. After the quartz capillary reached the desired depth, we applied silicone adhesive (KWIK-SIL, WPI) to seal the skull opening and fixed the quartz capillary position using dental cement (Contemporary Ortho-Jet™ Powder and Liquid, Lang Dental). Typically, the imaging experiment was performed ~4 weeks after this insertion process. To protect the capillary from dust, we put a plastic cap on the top of the head-bar, which was designed to match the cap diameter. It is worth noting that the 15 μm thin quartz capillary was strong enough to handle the brain motion. During the ~4 weeks of recovery, the mice were free to run, move and engage in routine activity. None of the implanted capillary was affected by these activities.

### Imaging probe bonding

The imaging probe is the combination of a high refractive index (N-LASF31) right-angle micro prism and a 1 mm diameter GRIN lens. To mount the probe onto the rotary stage, we fabricated a round holder (**Supplementary Fig. 7a**). To attach the probe onto the holder, we utilized a 2-axis translation stage to support the probe. We adjusted the position of the probe under a stereoscope such that its center was well aligned with the round opening on the holder. Then we pushed the translation stage along the cage rods so that the entrance facet of the GRIN lens went through the round opening (**Supplementary Fig. 7b**). We applied a tiny amount of UV epoxy to the gap between the GRIN lens and the round opening and cured the epoxy.

### Imaging probe alignment

To form a 360-degree panoramic view around the optical channel, we need to spin the imaging probe, which requires precision alignment of the probe such that its axis is aligned with the rotary stage axis. Using two orthogonal machine vision cameras (DMK 23UX249, The Imaging Source), we recorded 12 images of the probe on each camera during a 360-degree rotation controlled by the rotary stage (DT-80, PI). Using computation edge detection, we numerically determined the two edges of the probe. The average of the 24 edges from the 12 images revealed the true rotation axis of the rotary stage. With the assistance of these two cameras, we used micrometers to move the probe holder such that the probe’s axis was right on the rotation axis.

The capillary inside the mouse brain also needs to be aligned with the rotation axis. As the capillary is buried inside brain tissue, we utilized a downward imaging camera (camera 3 in **Supplementary Fig. 1**) equipped with a telecentric lens for the capillary alignment. An important step is that we need to transfer the previously measured rotation axis onto the imaging axis of camera 3. We first positioned a glass capillary within the view of camera 1 and 2. We oriented the capillary such that its axis overlapped with the rotation axis. Next, we translated the capillary along an optical stainless steel rail (x26-512, Newport) to be under camera 3. We adjusted the orientation of camera 3 such that its imaging axis was aligned with the capillary axis. Through this process, we mapped the rotation axis recorded by cameras 1 and 2 onto the imaging axis of camera 3. For daily *in vivo* imaging experiments, we just need to position the mouse under camera 3 and align the capillary to the imaging axis (**Supplementary Fig. 1c, 1d**). To insert the imaging probe inside the capillary, we utilized cameras 1 and 2 to center the opening of the capillary with respect to the rotation axis and moved the capillary upward along the axis using a motorized stage (VP-25XL-XYZR, Newport) that held the mouse head-bar.

### Alternative probe holding method

For imaging behaving animals with strong movement, we developed an alternative method for holding the imaging probe (**Supplementary Fig. 4**). Instead of firmly holding the probe, we utilized a docking mechanism. Experimentally, we simply attached an O-ring onto the probe at the desired location and dropped the probe inside the capillary. The friction of the O-ring was sufficient to hold the probe and allowed it to move with the capillary. To assist the releasing of the probe, we utilized a finite element analysis tool (Autodesk Inventor) to design and utilized wire electrical discharge machining to fabricate a flexure structure based probe holder (**Supplementary Fig. 4c**), which allowed holding and releasing the probe with an extremely compact frame.

### Optical system calibration

The propagation of light through the 1-pitch GRIN lens and the high refractive index prism all introduced spherical aberration. We conveniently utilized the SLM (X10468, Hamamatsu) in the imaging system to compensate for it. The most straightforward wavefront measurement is interferometry, especially off-axis holography^25^, which demands a long coherence length. We used an optical grating to couple the output of the femtosecond laser (Coherent Discovery) into a single mode fiber (**Supplementary Fig. 8a**). This step greatly reduced the spectral bandwidth and thus led to longer coherence length needed for the off-axis holography. First, we utilized off-axis holography to record the phase profile at the pupil plane (**Supplementary Fig. 8b**). Next, we commanded the SLM pixels to change phase. Through this process, we determined the relationship between the SLM’s pixel and the hologram recording camera’s pixel. We then displayed the reversed phase on the SLM to compensate for the aberration. The aberration of the reference beam is not negligible. To fully eliminate the residual aberrations, we utilized a self-reference measurement method (**Supplementary Fig. 8c**), named IMPACT^26^, for wavefront measurement (**Supplementary Fig. 8d**). Finally, the combined wavefront (**Supplementary Fig. 8e**) was applied to the SLM to compensate for the system aberration.

## Acknowledgement

This work was funded by NIH (U01NS094341, U01NS107689) and Purdue University. M.C. thanks Dr. Scott Sternson of HHMI Janelia research campus and Prof. Jeffry Isaacson of UCSD for valuable comments and discussion, and thanks Howard Hughes Medical Institute for scientific instruments.

## Author Contributions

M.C. invented COMPACT, designed and implemented the imaging system. M.C. and W.G. supervised the research. W.G. designed the biology experiment. B.W. C.W. and Z.C. collaborated on the experiment, data analysis, and figure preparation. All authors contributed to the writing of the manuscript.

## Correspondence

Correspondence and requests for materials should be addressed to M.C.

## Supplementary Material

**This file includes Supplementary Figure 1-8 and caption for video 1-5.**

**Supplementary Figure 1.**
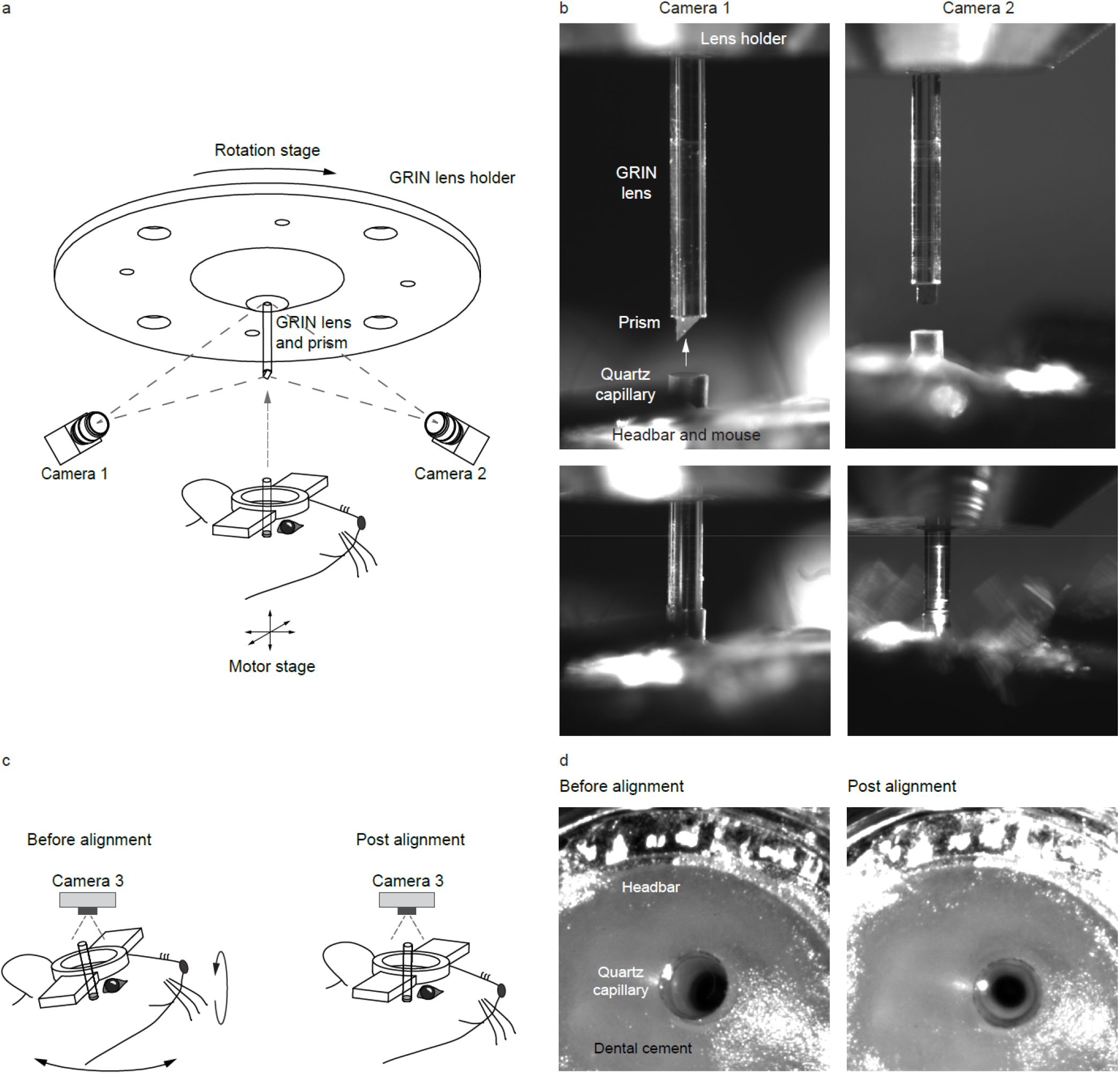
Procedure for imaging probe alignment. (a) Imaging probe alignment. As the imaging probe need to spin inside the capillary, both the imaging probe and the capillary need to be well aligned with the rotation axis of the rotary stage. We commanded the rotary stage to spin in 12 steps over 360 degrees, during which we used two machine vision cameras (camera 1 and camera 2) to record the imaging probe position. From each image, we obtained the boundary position of the probe. The average of the 24 boundaries yielded the rotation axis. With the aid of the two machine vision cameras, we aligned the imaging probe to the rotation axis. (b) Representative images from camera 1 and 2. (c) Capillary alignment. As the capillary was mostly embedded inside the brain tissue, viewing from the top is much more convenient than using the side viewing cameras. To assist the daily routine alignment, we set up a third machine vision camera (camera 3) equipped with a telecentric imaging lens. The imaging axis of the telecentric lens was aligned with respect to the rotation axis. (d) For routine alignment, we just need to view the capillary from the top using camera 3 and align the capillary to the imaging axis.

**Supplementary Figure 2.**
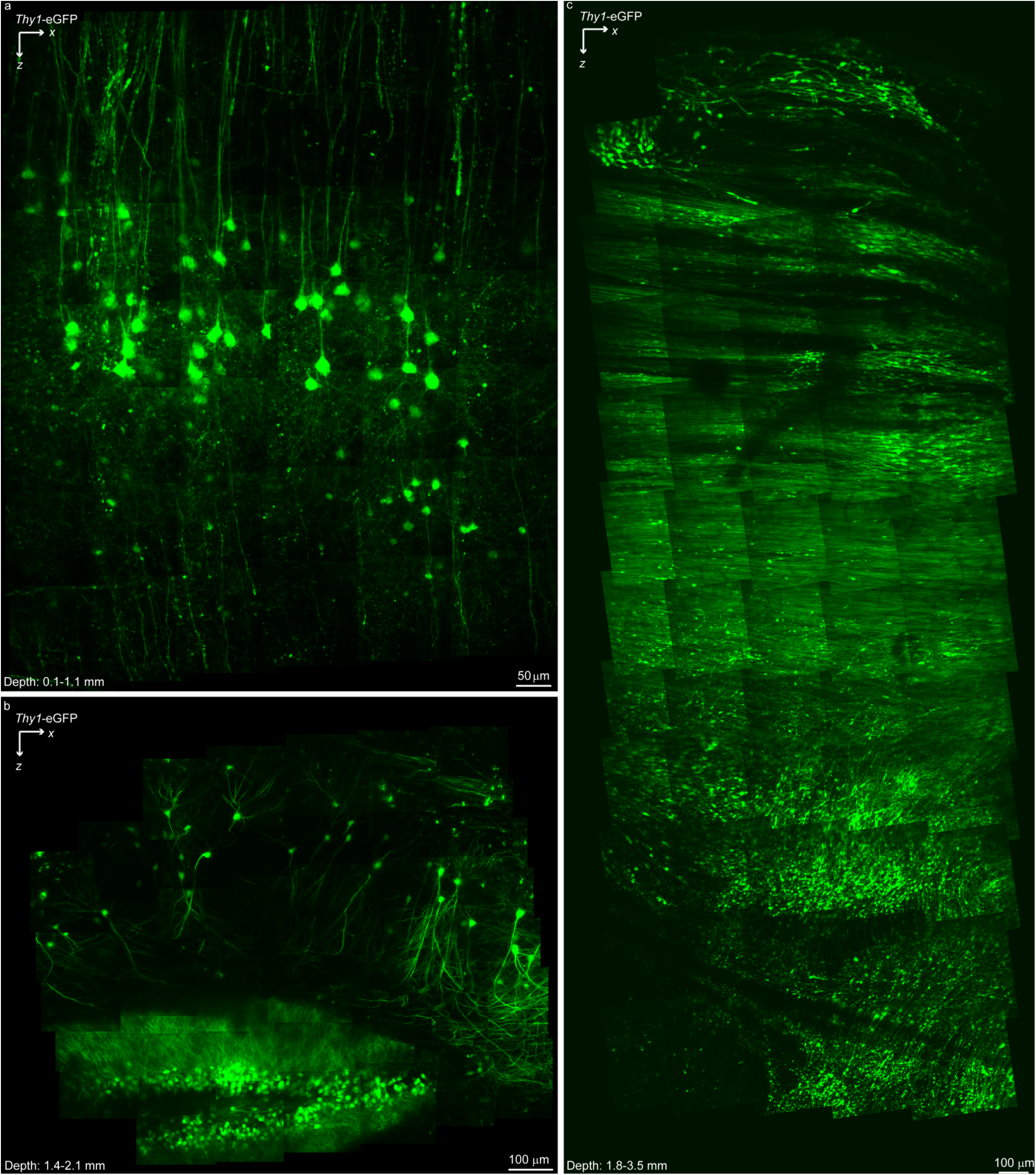
*In vivo* two-photon imaging of neuronal structure in eGFP expressing mouse brain at different depths. (a) Imaging neocortex from 0.1 to 1.1 mm. The dendrites and axons can be clearly seen. (b) Imaging hippocampus from 1.4 to 2.1 mm. (c) Imaging thalamus from 1.8 to 3.5 mm.

**Supplementary Figure 3.**
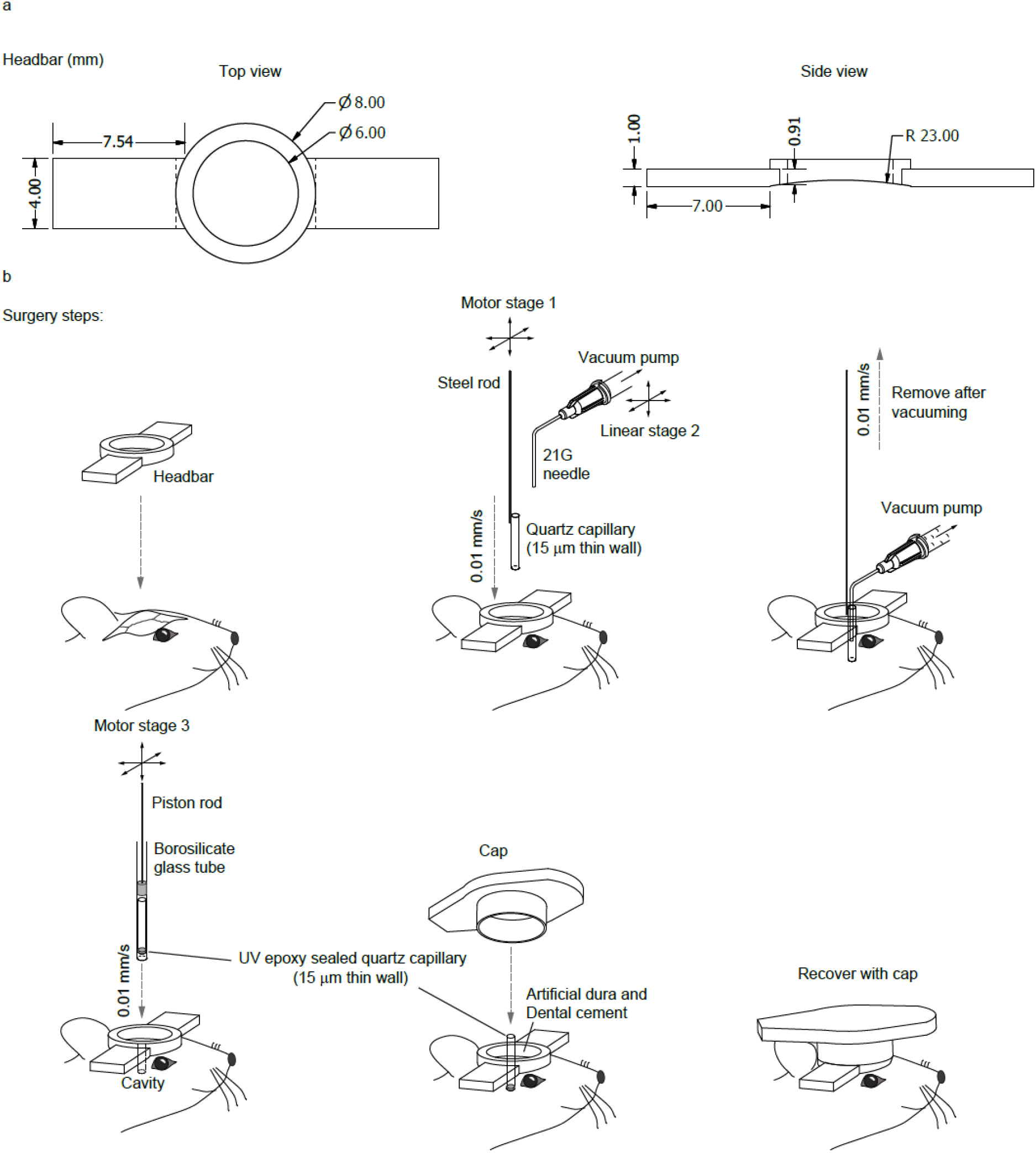
Surgical procedure for capillary implantation. (a) Design of the head-bar for head fixed animal imaging. The ring is slightly above the bar to couple with the cap that will be applied at the end of the surgery. (b) Surgical procedures. We first removed the skin above the skull and attached the head-bar to the skull. Next, we opened a 3 mm diameter aperture on the skull and used a computer controlled actuator to gradually insert an open-end quartz capillary at 10 μm/sec speed. During the process, we used a 21G needle connected to a vacuum pump to remove the tissue enclosed by the capillary. After reaching the desired depth, we removed the open-end capillary and injected the closed-end capillary inside the brain tissue. Essentially, we employed the open-end capillary to first create an opening in the tissue and then filled the opening with the closed-end capillary (the imaging channel). To guide the capillary straight into the tissue, we loaded the capillary inside an injection tube formed by the glass tube and plastic piston used in transfer pipettes. A motorized actuator slowly pushed the piston at 10 μm/sec speed. Finally, we put a plastic cap on top of the head-bar to protect the capillary from dust.

**Supplementary Figure 4.**
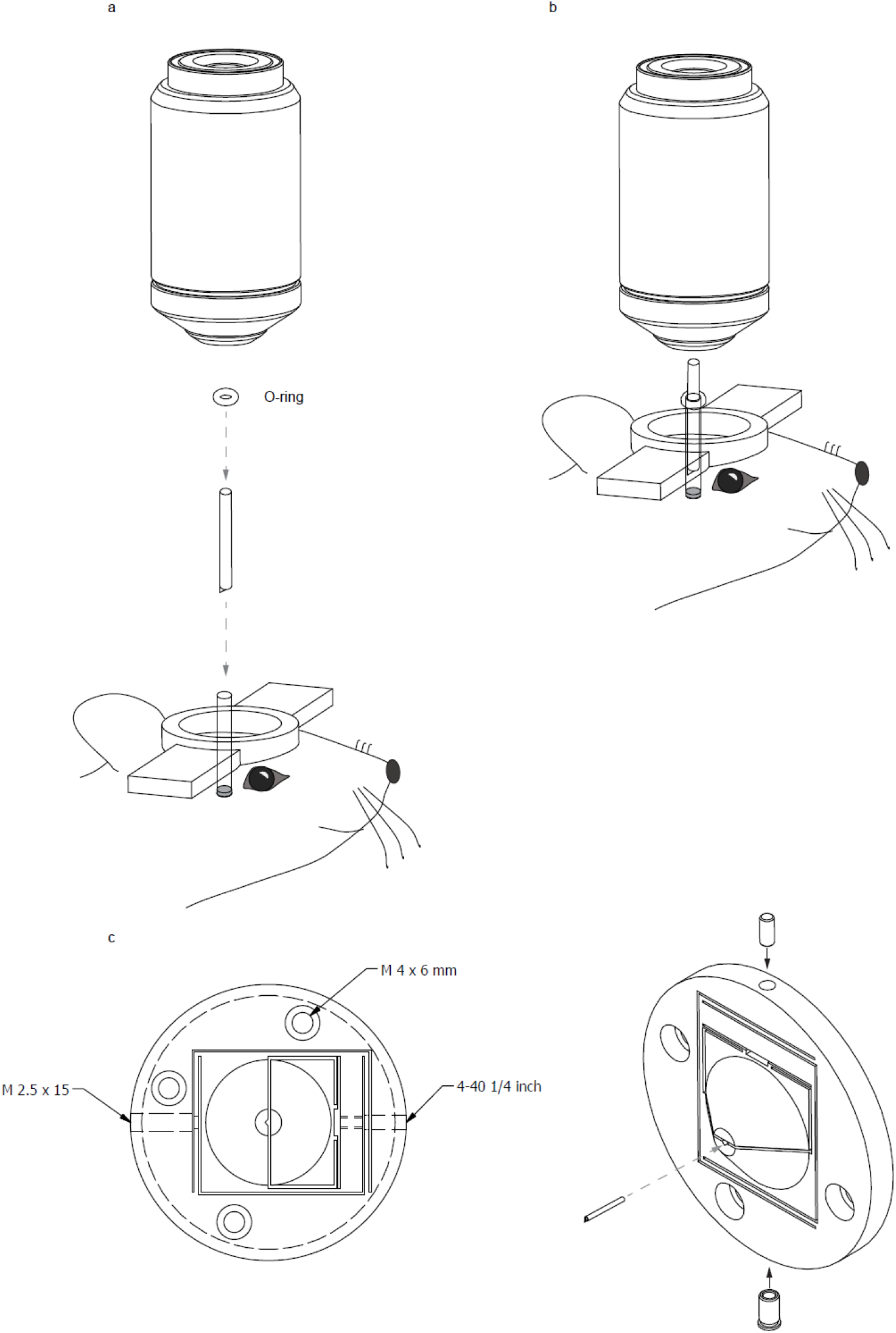
Imaging probe docking method. (a) Instead of holding the imaging probe with the rotary stage, we attached a O-ring to the probe at the desired location and released the probe into the capillary. The friction of the O-ring was sufficient to hold the probe. (b) With the O-ring based docking, we can directly position the mouse under the objective lens for two-photon calcium imaging. (c) For finer adjustment on probe orientation and depth, we designed and fabricated a flexure structure based quick-release probe holder. There are two fine screws on the flexure holder. One is used for locking the probe and the other is used to fine tune the position of the probe. With this flexure holder, we can release the imaging probe at the desired orientation and depth.

**Supplementary Figure 5.**
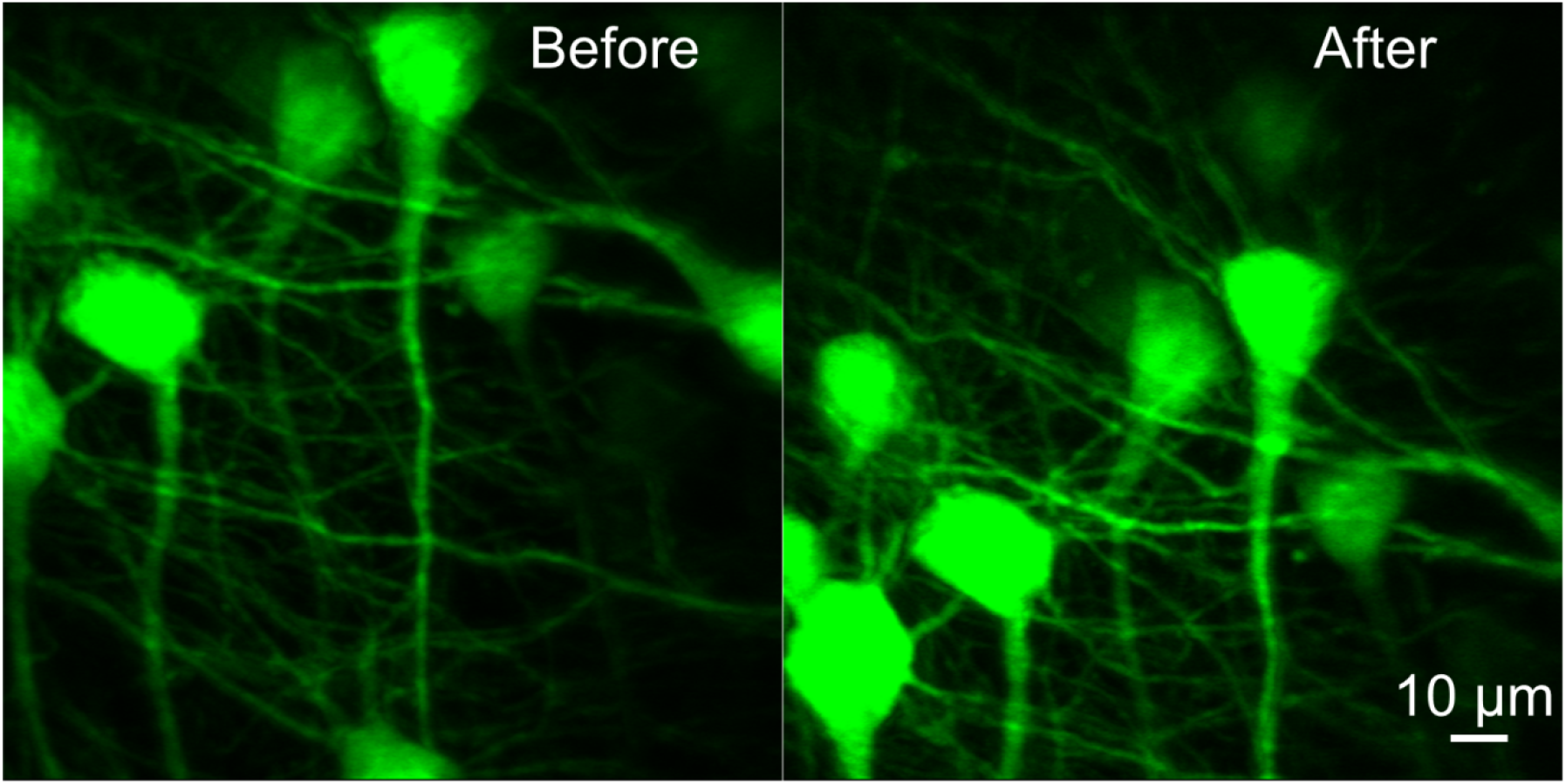
Probe insertion repeatability test. We reloaded the same eGFP mouse to the imaging system and commanded the COMPACT system to image the same region (same insertion length, angle, and imaging depth). The same group of neurons can be conveniently relocated. The position offset between the reloading was within tens of microns. By using the machine vision camera to reset position 0 based on the capillary opening position, we can potentially reduce the reloading accuracy to ~10 micron.

**Supplementary Figure 6.**
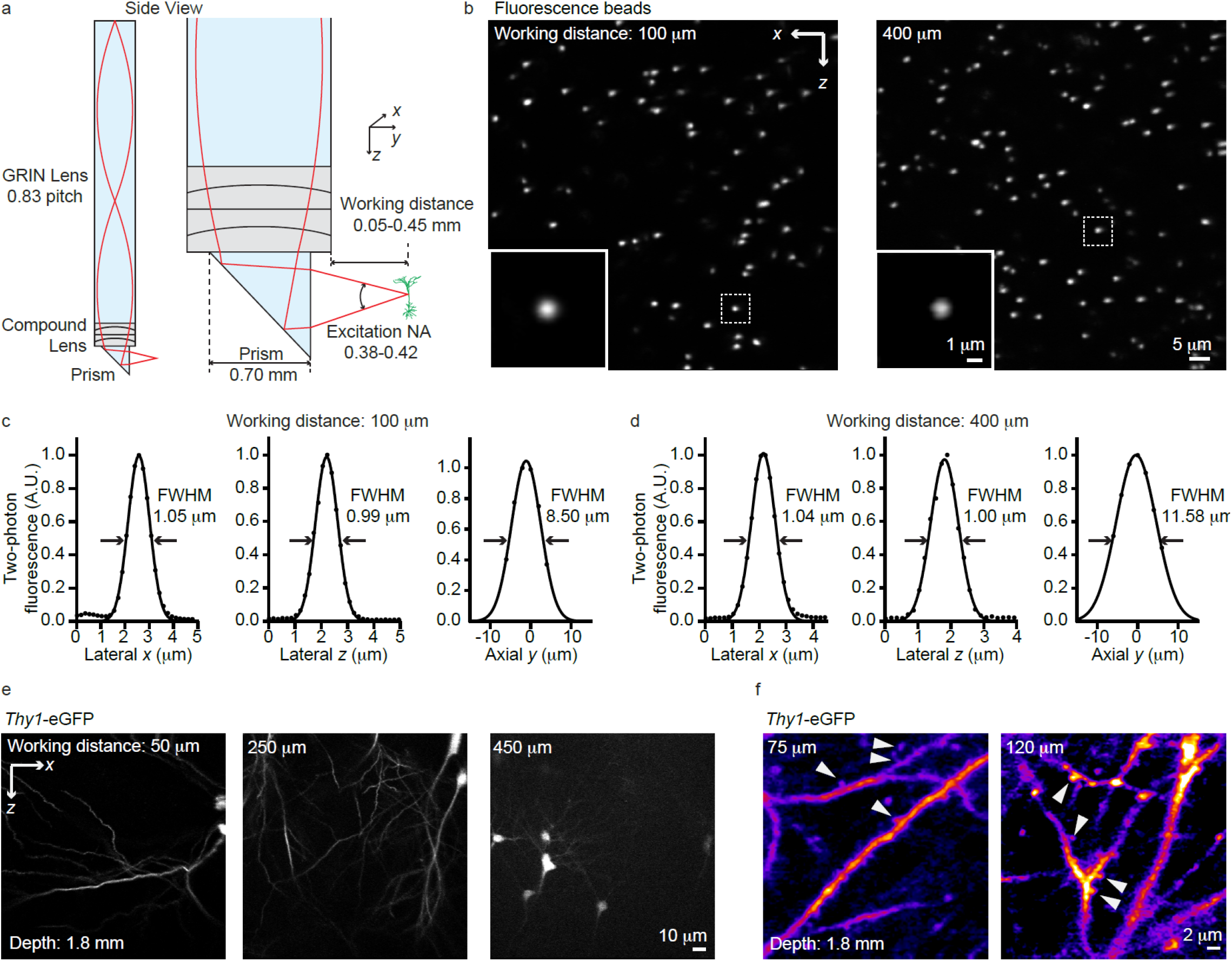
High-resolution compound probe for two-photon COMPACT imaging. Using compound lenses, we could increase the excitation NA for high-resolution imaging. (a) Compound lens configuration. The excitation NA reached 0.38-0.42 for imaging 50-450 μm outside the capillary. (b) Two-photon imaging of 0.5 μm fluorescence beads. (c, d) The imaging point spread function for imaging 100 and 400 μm outside the capillary, respectively. The transverse and axial resolutions were both improved compared to the simple probe that comprised only a GRIN lens and a prism mirror. (e) Two-photon imaging of eGFP mouse brain with an imaging depth of 1.8 mm. The maximum working distance reached 450 μm. (f) At shorter working distances, the fine neuronal structures such as the dendritic spines became visible. These results highlight the resolution advantage of the compound imaging probe.

**Supplementary Figure 7.**
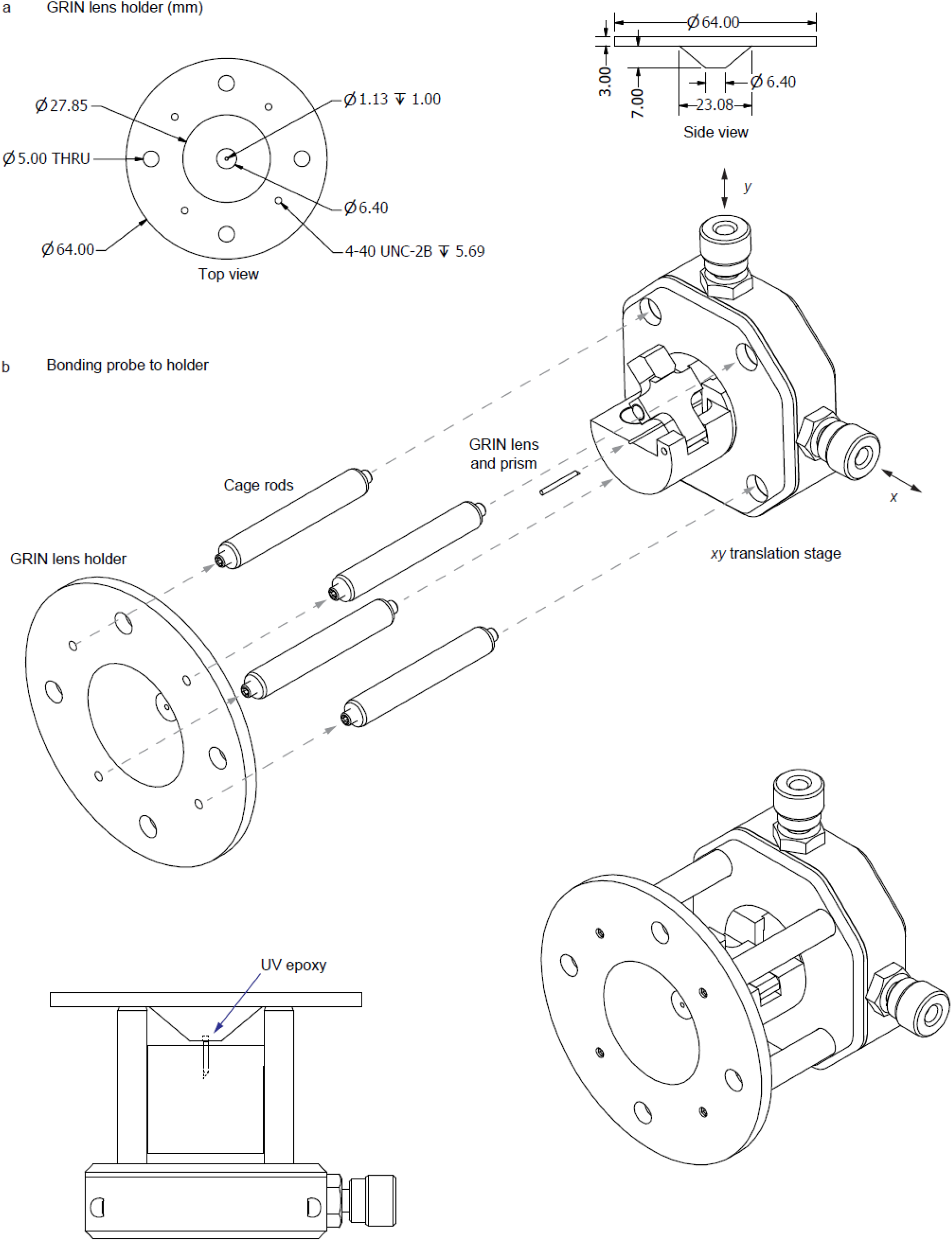
Procedure for attaching the imaging probe onto its holder. (a) The dimension of the imaging probe holder. Units: mm. (b) Alignment device which utilized the cage system from Thorlabs. We observed the imaging probe through the central aperture of the holder under a stereoscope and utilized a 2-axis translation stage to center the probe on the aperture. A light amount of UV epoxy was applied to the gap between the imaging probe and the holder, followed by UV curing.

**Supplementary Figure 8.**
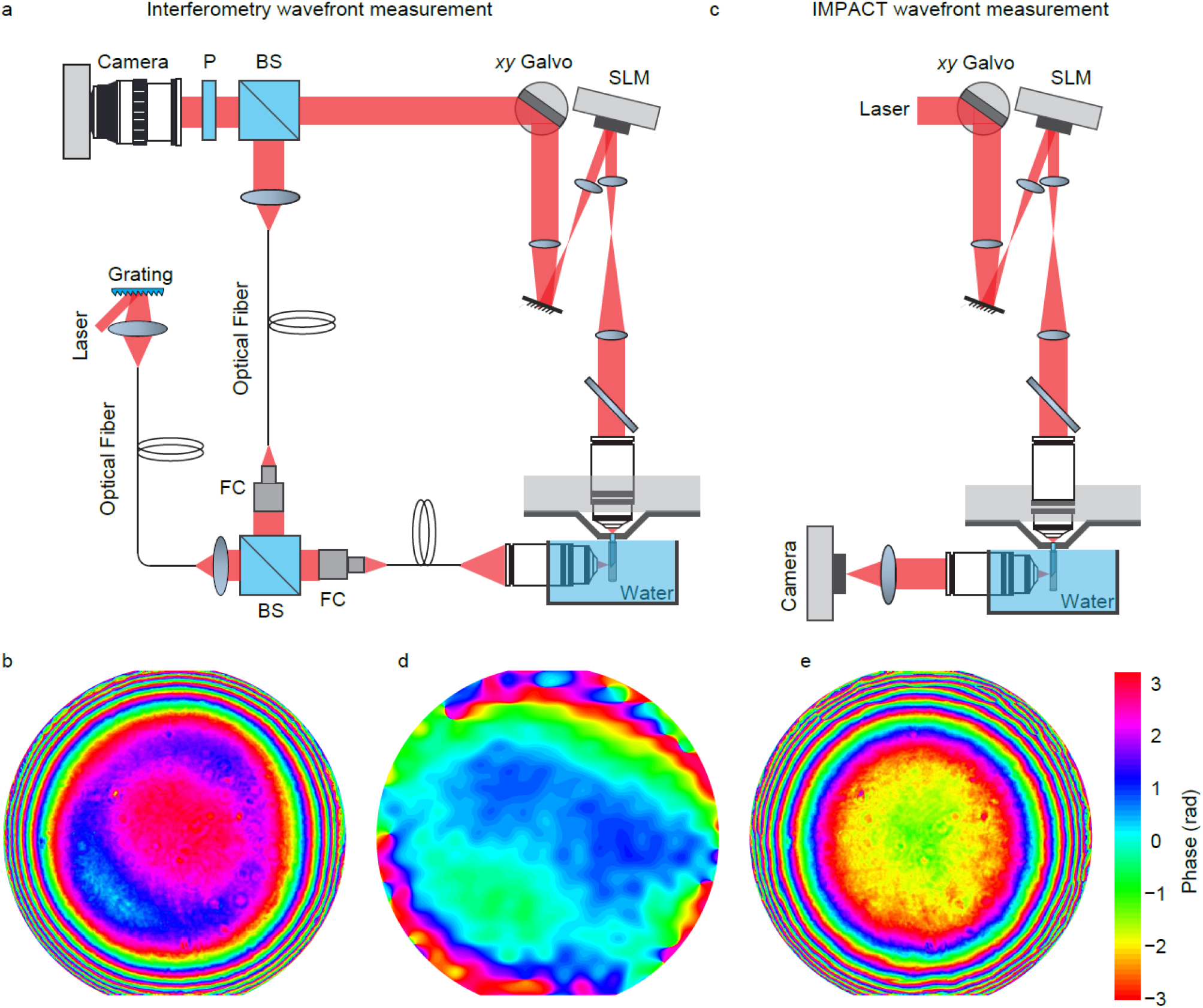
Imaging system optical calibration. (a) The optical design for single-shot off-axis holography measurement of the imaging system aberration. The laser source was the actual two-photon imaging laser. The grating coupled single mode fiber light delivery greatly reduced the bandwidth and thus reduced the dispersion and increased the coherence length. (b) The off-axis holography measured system aberration, which was dominated by the spherical aberration induced by the imaging probe. (c) To account for the residual aberration in the off-axis holography measurement, we utilized a modulation based wavefront correction method, named IMPACT, to optimize the focus peak intensity. (d) The residual wavefront measured by IMPACT. (e) The combined wavefront was displayed on the SLM to compensate the system aberration.

## CAPTIONS FOR SUPPLEMENTARY VIDEOS

**Supplementary Video 1.**
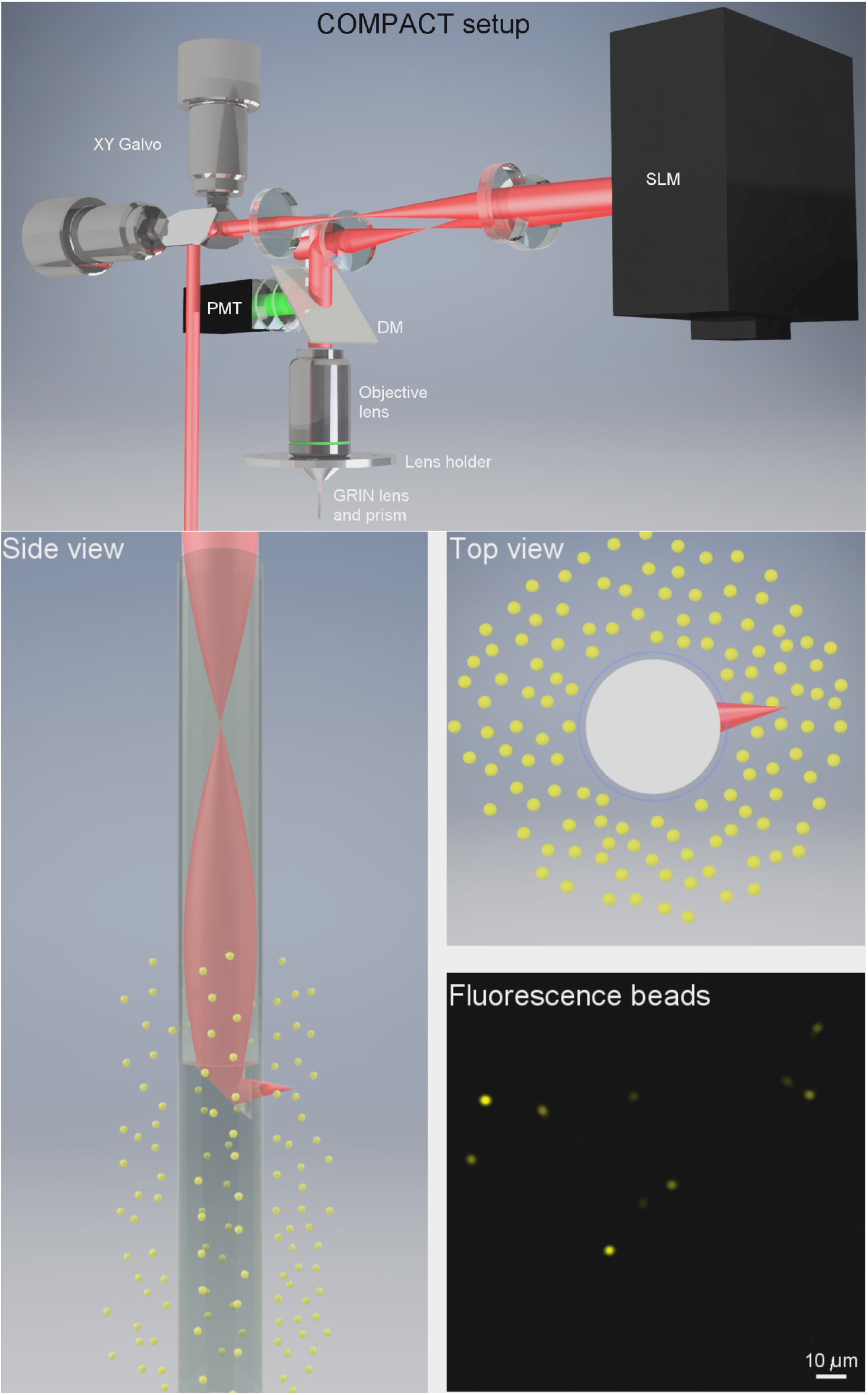
Design of the two-photon COMPACT imaging system. The optical configuration of the two-photon imaging system and the operation of whole-depth panoramic two-photon imaging.

**Supplementary Video 2.**
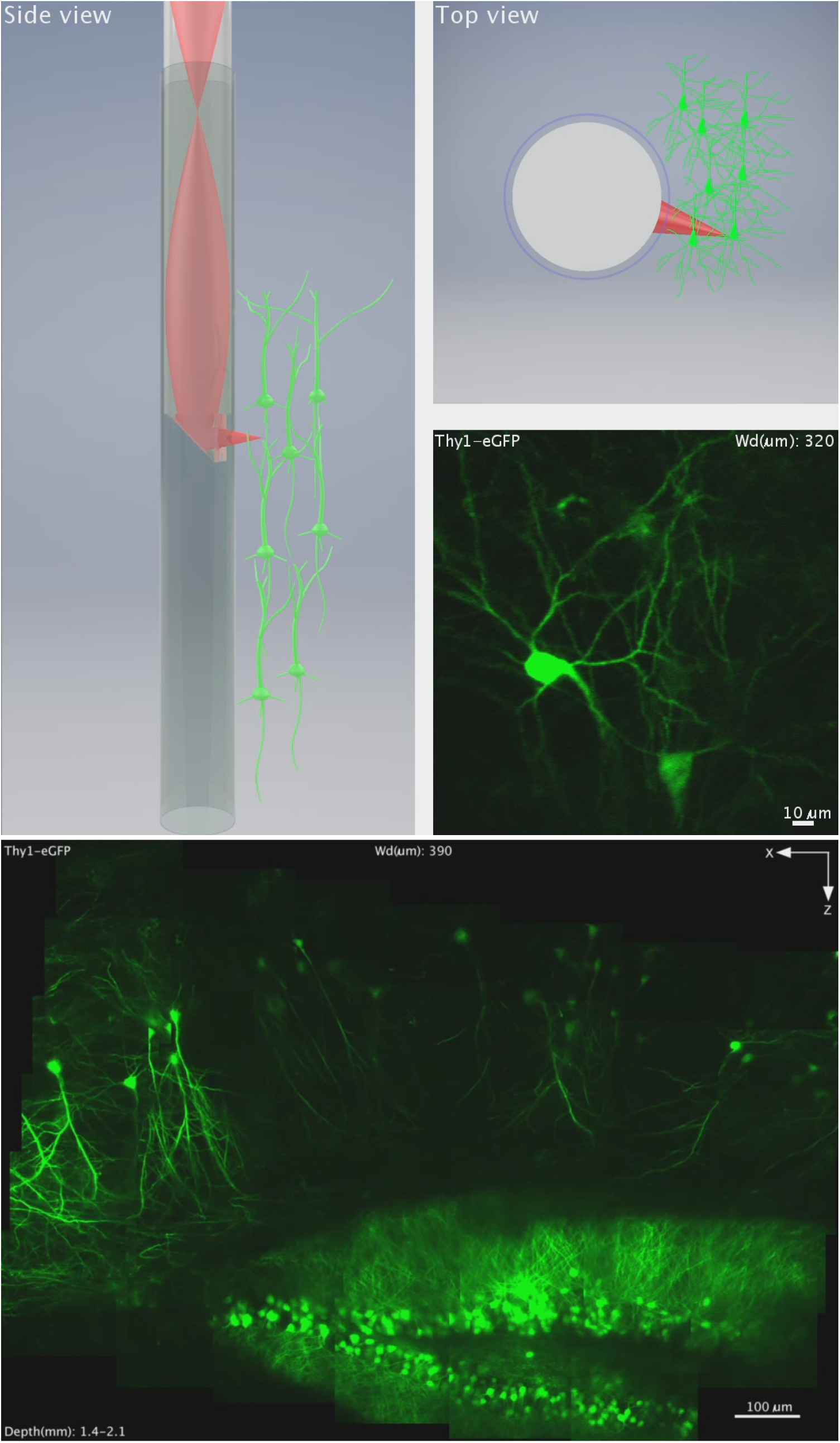
Two-photon structural imaging of eGFP mouse brain. The acquisition of the image volume shown in **Supplementary Fig. 2b**.

**Supplementary Video 3.**
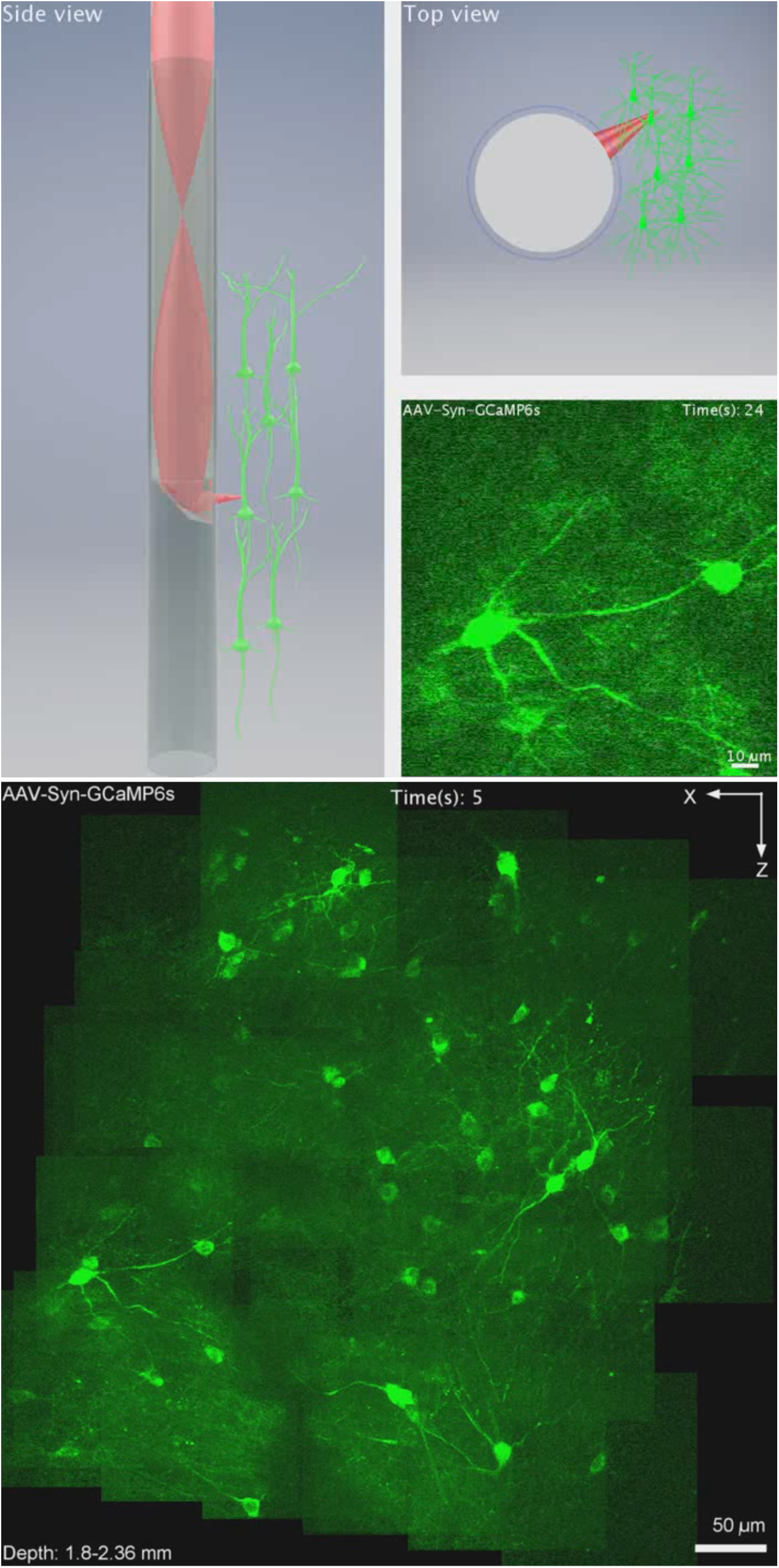
Two-photon calcium imaging of awake mouse brain. The acquisition of the image volume shown in **Fig. 3**.

**Supplementary Video 4.**
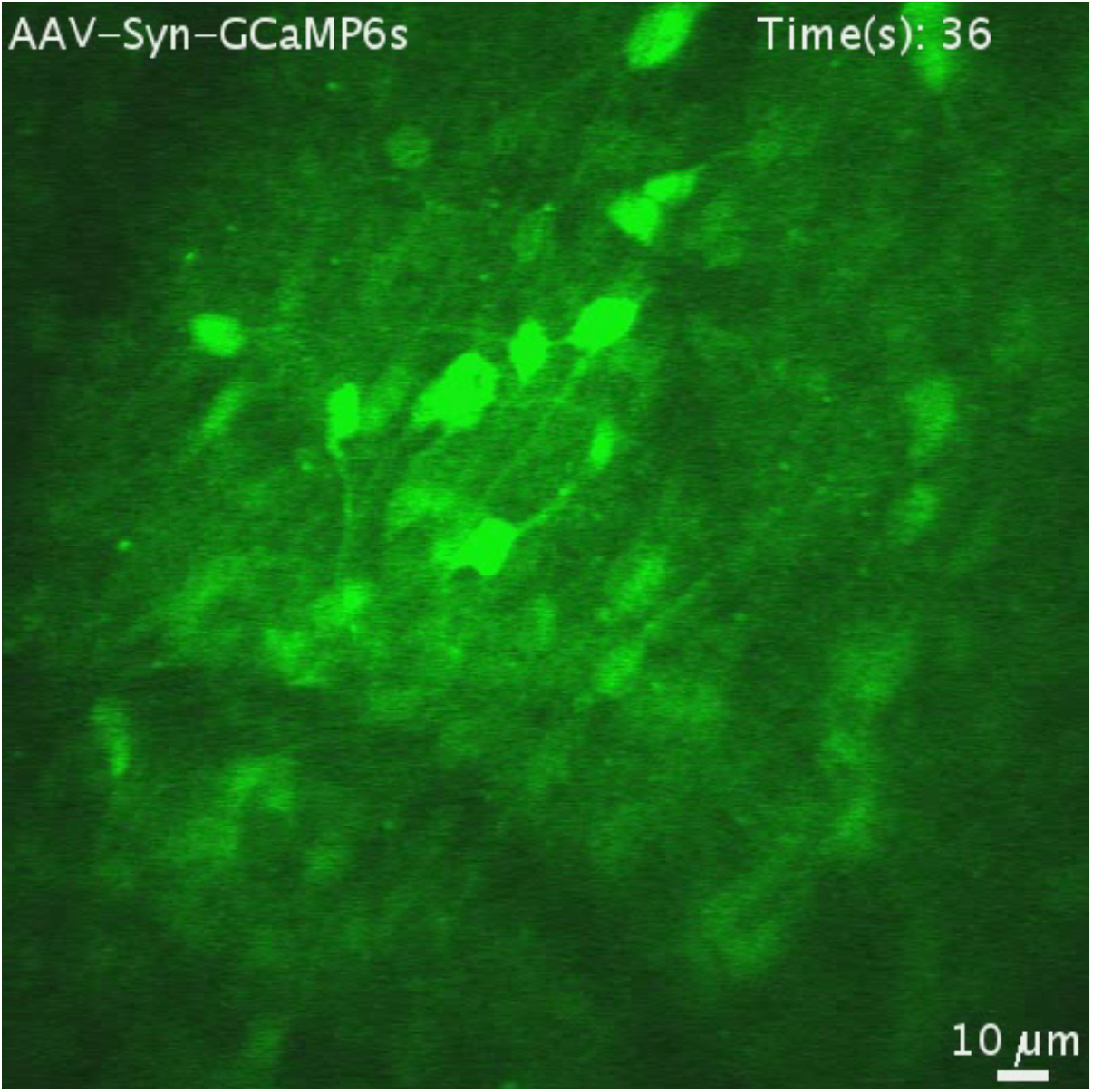
Two-photon calcium imaging of awake mouse brain using the probe docking method.

**Supplementary Video 5.**
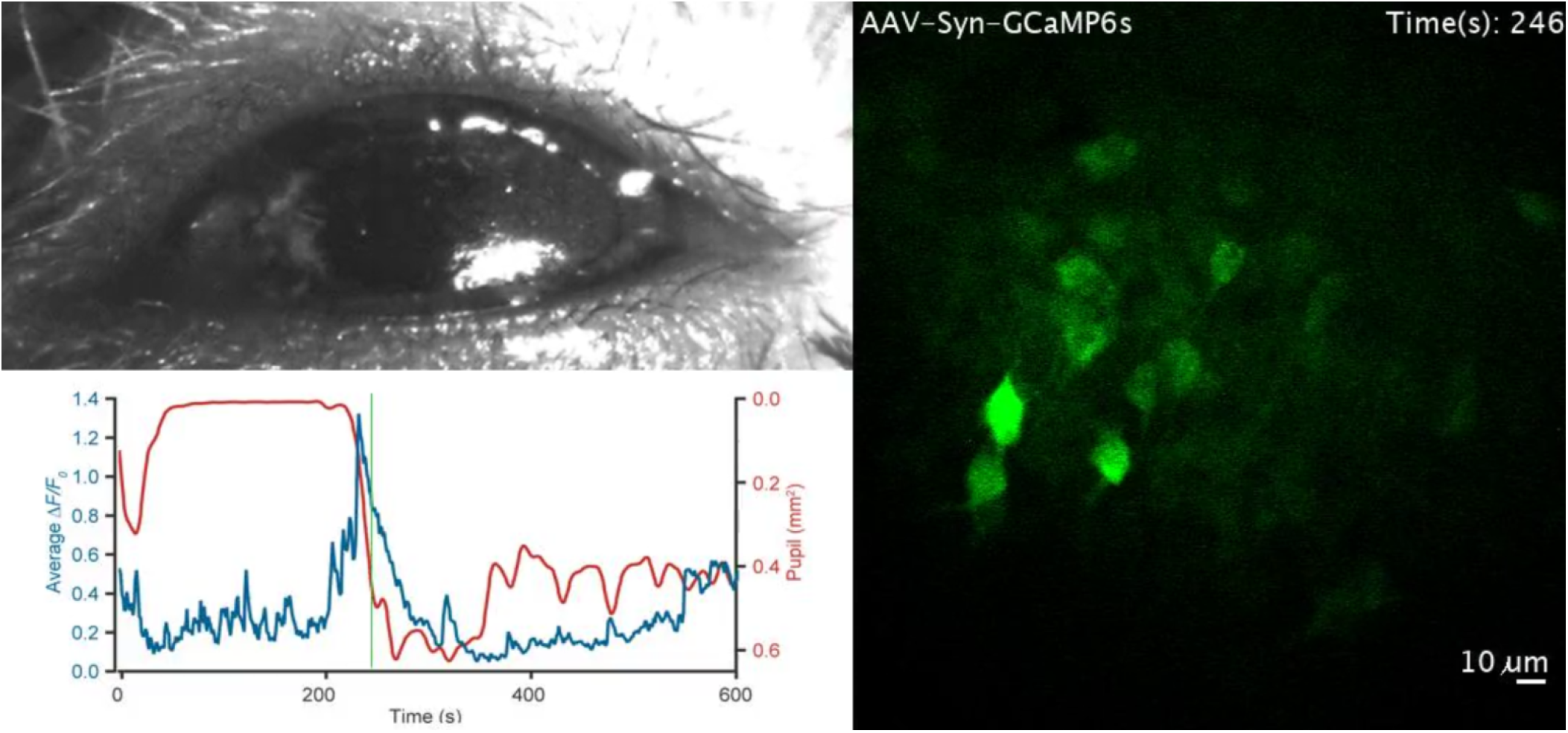
Longitudinal multiregional calcium imaging of neuronal activity associated with sleep.

